# Collateral sensitivity increases the efficacy of a rationally designed bacteriophage combination to control *Salmonella enterica*

**DOI:** 10.1101/2023.09.19.558423

**Authors:** Luke Acton, Hannah Pye, Gaëtan Thilliez, Rafał Kolenda, Michaela Matthews, A. Keith Turner, Muhammad Yasir, Emma Holden, Haider Al-Khanaq, Mark Webber, Evelien M Adriaenssens, Robert A Kingsley

**Affiliations:** Quadram Institute Biosciences, Norwich Research Park, Norwich, NR4 7UQ, UK; University of East Anglia, Norwich, NR4 7TJ, UK

**Keywords:** Host-Range, Collateral Sensitivity, TraDIS, LPS, BtuB, Phage Cocktail

## Abstract

The ability of virulent bacteriophages to lyse bacteria influences bacterial evolution, fitness, and population structure. Knowledge of both host susceptibility and resistance factors is crucial for the successful application of bacteriophages as biological control agents in clinical therapy, food processing and agriculture. In this study, we isolated twelve bacteriophages termed SPLA phage which infect the foodborne pathogen *Salmonella enterica*. To determine phage host range, a diverse collection of *Enterobacteriaceae* and *Salmonella enterica* were used and genes involved in infection by six SPLA phages were identified using *Salmonella* Typhimurium strain ST4/74. Candidate host receptors included lipopolysaccharide, cellulose and BtuB. Lipopolysaccharide was identified as susceptibility factor for phage SPLA1a and mutations in LPS biosynthesis genes spontaneously emerged during culture with *S*. Typhimurium. Conversely, LPS was a resistance factor for phage SPLA5b that suggested that emergence of LPS mutations in culture with SPLA1a may represent a case of collateral sensitivity to SPLA5b. We show that combination therapy with SPLA1a and SPLA5b was more successful in limiting the emergence of phage resistance compared to monotherapy. Identification of host susceptibility and resistance genes and understanding infection dynamics is critical step in rationale design of phage cocktails against specific bacterial pathogens.

## Introduction

Viruses of bacteria (phages) infect and lyse bacteria and as such have the potential to replace or complement the use of antibiotics to treat bacterial infections or to control bacterial pathogens in the environment (1–3). The emergence of difficult to treat, antibiotic resistant bacterial infections has renewed interest in developing products using lytic phages. Phages initiate infection of the bacterial host by specific binding to a receptor or receptors located on the cell surface of a susceptible bacteria. In Gram-negative bacteria such as *Salmonella enterica*, well characterised receptors include outer membrane proteins (OMPs), lipopolysaccharide (LPS) and flagella antigens (4). Receptor specificity of phage infection presents two challenges for the implementation of phage as antimicrobials, host specificity that limits spectrum of activity, and the potential for emergence of resistance through a single mutation. To overcome these limitations, phages are commonly pooled together to form a phage cocktail that both extends host range and limits the emergence of resistance. Selection of phages to be included in a phage cocktail should be based on a robust understanding of the bacteriophage host range, targeted receptor and mechanisms of host resistance (5). However, mechanistic data for phage activity is often limited resulting in suboptimal cocktail design and failed applications (6, 7).

Non-Typhoidal *Salmonella* (NTS) remains an important pathogen which poses a significant threat to human and livestock health, and the economy. NTS is estimated to cause 93.8 million diarrheal illnesses and cause 155,000 deaths each year (8), with an associated economic cost estimated to exceed 3 billion Euros in the European Union alone (9). Most NTS infections result from the consumption of food or water that has become contaminated by faeces of infected animals. Control of *Salmonella* in livestock has become increasingly challenging as resistance to commonly used veterinary antibiotics has increased and concerns over the use of veterinary antibiotics leading to resistance to related antibiotics used to manage clinical infections in people, has limited their use. Bacteriophages are a particularly promising solution to control *Salmonella* in livestock (7), but may also be appropriate in the environment of the post-slaughter food chain where antibiotics use is not possible.

We report the isolation of twelve *Salmonella enterica* bacteriophages with diverse host- range and host susceptibility and resistance genes. Prediction of potential mechanisms of resistance using data from a transposon insertion mutant library functional genomics screen enabled rationale design of a combination of phage that exploited collateral sensitivity that resulted from resistance to one phage to increase efficacy.

## Materials and Methods

### Selection of Bacterial Strains

Twelve *Salmonella enterica* s t r a i n s w e r e u s e d f o r t h e enrichment of environmental samples for bacteriophages (Supplementary Table 1). Strains were of serovar Montevideo, Panama, Mbandaka, Kedougou, Infantis, Derby, Newport, Enteritidis and Typhimurium. Eleven strains were previously isolated from food products in the UK. The remaining strain was an isogenic mutant of *Salmonella enterica sv* Typhimurium (*S.* Typhimurium) strain ST4/74 that lacked the Gifsy-1, Gifsy-2, ST46B, SopEΦ and P4-like prophages generated previously (10). Strains used for analysis of bacteriophage host range were termed HRS (Host Range Strain). This collection comprised of thirty-six strains including *Salmonella enterica* from various sources, a s w e l l a s o t h e r b a c t e r i a l s p e c i e s isolated from food products (Supplementary table 1).

### Bacterial Culture and Allelic Exchange

Strains were routinely cultured in Lysogenic Broth (LB) or on LB containing 1.5% agar. Where appropriate, cultures were supplemented with kanamycin (50 µg/ml) or hygromycin (75µg/ml). Construction of mutant strains, with the exception of *S*. Typhimurium ST4/74 Δ *rfaK,* was performed by one-step inactivation using a method described previously (11) but using the pSIM18 plasmid (12). PCR products were generated using primers listed (Supplementary Table 2). The primers amplified the *aphII* gene from plasmid pKD4 and tagged it with 50bp regions which were homologous to the target for mutagenesis at both ends. Allelic exchange was performed in *S.* Typhimurium ST4/74 containing pSIM18 (13). *S*. Typhimurium ST4/74 Δ *rfaK* was generated using gene doctoring using plasmids previously designed by Thompson *et al* (14).

### Isolation of Bacteriophages

All environmental samples were collected between November 2019 and February 2020. The collection included eight wastewater treatment samples, eight retail meat samples, nine lake/river samples and six samples from drains located in a food production factory. Samples from food were enriched in buffered peptone water (BPW) by incubation at 37⁰C for 18 hours with shaking. To isolate bacteriophages, 25ml of sample was centrifuged at 3220 x g for 10 minutes. The supernatant was filtered through a 0.45µM pore size, polyethersulfone (PES) sterile syringe filter and 5ml added to 40ml of 2x LB Broth along with 200 µl of a mixture of bacterial strains cultured to mid exponential phase (OD600_nm_ of 0.6). The sample and culture were incubated at 37 ⁰C for 18 hours. The resulting culture was centrifuged and filtered as previously described. New phage isolates were identified by spot assay on double agar overlay containing each of the enrichment strains to identify the host *S. enterica* strain with the greatest sensitivity to lysis as described previously (15). All phage preparations were confirmed to contain a single DNA bacteriophage based on assembly of short reads into a single contiguous sequence and a single plaque morphology in overlay assay.

### Phage and bacterial genomic DNA Extraction and Sequencing

For preparation of phage genomic DNA, high titre lysates (>10 ^9^ PFU/ml) were prepared by enrichment in broth cultures of the host strain that exhibited the clearest plaques and were used for nucleic acid extraction. A 1 ml aliquot of each phage was treated with 1 µl DNase (New England Biolabs, USA) at 37 ⁰C for 40 minutes. Phage virions were concentrated with 500ul PEG solution (24% Polyethylene glycol mw 8000, 1M NaCl) overnight at 4 ⁰C and subsequently resuspended in 200 µl of nuclease free water. Nucleic Acid Purification with Proteinase K digestion was performed using Maxwell ® RSC Total Viral Nucleic Acid Purification Kit according to the manufacturer’s protocols (Promega, USA). For preparation of bacterial genomic DNA, strains were cultured at 37⁰C for 18 hours in LB broth with shaking, DNA was extracted using Maxwell® RSC Cultured Cells DNA kit (Promega, USA) and NexteraXT sequencing libraries were prepared following the manufacturer’s protocols (Illumina, USA) and sequenced using the Illumina NextSeq500 platform.

### Bioinformatic Analysis

Quality control of raw, paired end, Illumina sequencing reads carried out was using fastP (v0.23.2) with default parameters (16). Reads passing quality control were assembled *de novo* using the SPAdes based assembler Shovill (v1.0.4) (17, 18). For bacteriophages, sequence reads were aligned to assembled phage contigs using BWA-mem (v0.7.17.1) (19). A tree based on sequence similarity of the proteome sequence was calculated using tBLASTx within the VIPtree software package and annotated using the INPHARED database (20, 21). Average nucleotide identity (ANI) comparisons were made using FastANI (v2.0) (22) . For phylogenetic reconstruction of bacterial strains of multiple genera of the order Enterobacterales, paired end reads were assembled *de novo* using the SPAdes based assembler Shovill (v1.0.4) (17, 18) and the 16S rRNA gene identified using barrnap (https://github.com/tseemann/barrnap). Multiple sequence alignments of the 16S rRNA gene was performed using MAFFT (v7.470) (23) and the alignment trimmed using TrimAL (24). The resulting 1536 base pair alignment was used to generate a maximum likelihood phylogenetic tree using IQtree 2 (v2.1.4-beta) with model selection with ModelFinder (25, 26). The selected substitution model was HKY+F+R2 with an ultrafast bootstrap value of 1000 (27). For phylogenetic construction of *Salmonella* strains, paired end sequences were aligned to the *S*. Typhimurium SL1344 reference genome (FQ312003) using the rapid haploid variant calling and core SNP phylogeny pipeline SNIPPY (https://github.com/tseemann/snippy) (v4.3.6) . Maximum likelihood phylogenetic trees were constructed using the multiple sequence alignment with RAxML (v8.2.12) (28) using the GTRCAT model with a bootstrapping value of 100.

### Bacteriophage host range, growth curve analysis and liquid assay score (LAS)

Bacterial strains were cultured in LB broth for 18 hours at 37 ⁰C at 200rpm, adjusted to 1 x 10 ^7^ CFU/ml and 180 µl added to each well of a 96-well Cyto-One microtiter plate (Starlab, UK). Bacteriophages were inoculated into wells at a multiplicity of infection (MOI) of 1. Plates were placed into FLUOstar Omega Microplate reader (BMG LABTECH, Germany) incubated at 37 ⁰C with double orbital shaking and the optical density (600nm) was measured every 15 minutes for 18 hours. Bacterial growth curves were plotted using mean baseline corrected Optical Density readings against time with 3 biological and technical replicates. Area under the curve (AUC) was calculated using the previously published method by Xie et al (29). Frequency of resistance to phages SPLA1a and SPLA5b in monotherapy and combination therapy was assessed using the same method with 5 biological and 96 technical replicates. The difference in AUC (liquid assay score) was calculated using Equations 1 and 2.

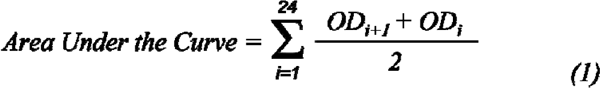

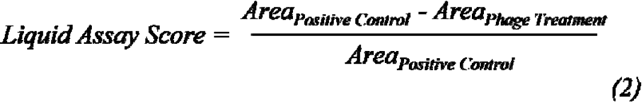

***Equation 1-2 – Method used to quantify growth curve data into liquid assay scores (LAS) scores***.

### Construction of transposon mutant library

To construct a Tn5 transposon mutant insertion library in *S*. Typhimurium ST4/74, transposon DNA was amplified by polymerase chain reaction (PCR) using P-Tn5Km-01 and P-Tn5Cm-04 oligonucleotides (sequences available in supplementary table 2) and a custom plasmid, pHPTTn5Km, as template DNA (Supplementary figure 1). PCR products were purified using QIAquick PCR Purification Kit (Qiagen, Germany). Transposomes were prepared using 100 ng transposon DNA and EZ-Tn5 transposase (Lucigen, USA) according to the suppliers specifications. For preparation of electrocompetent cells, a culture of *S*. Typhimurium ST4/74 was first prepared in 5mL LB broth and incubated for 18 hours at 37°C in a shaking incubator (200rpm). A 500µL aliquot of this culture was added to 50mL 2xYT broth (1.6% tryptone, 1% yeast extract, 85.6 mM NaCl) and incubated at 37°C to OD _600nm_ of 0.2-0.25. Cells at the required optical density were harvested by centrifugation at 3500g for 10 minutes at 4°C. The resulting cells were washed three times with 10% glycerol and resuspended in 600 µl 10% glycerol. A 60 µL aliquot of electrocompetent bacterial cells were mixed with 2 µL sterile nuclease free water, 2 µL TypeOne Restriction Inhibitor (Lucigen, USA) and 0.4 µL transposome, on ice. Following electroporation, cells were immediately resuspended in 1 mL S.O.C (2% tryptone, 0.5% yeast extract, 10mM NaCl, 2.5 mM KCl, 10 mM MgCl _2,_ 10 mM MgCSO _4_ and 20mM glucose) prewarmed to 37°C) and recovered at 37 °C for 1.5 hours. The cell suspensions in S.O.C were then spread on LB agar supplemented with kanamycin at 50µg/mL and these incubated at 37°C for 16 h to select for transposon mutant colonies. Separate 10 µL and 100 µL volumes of the cell suspensions in S.O.C. were also spread on LB agar supplemented with kanamycin (50µg/mL) for enumeration to allow an estimation of the total number of mutants obtained. Resulting colonies were harvested into LB-broth and glycerol was added to a final concentration of 15%, then aliquots of this were stored at -80°C. For each experiment, a 50µL aliquot of this stored *S*. Typhimurium ST4/74 transposon mutant library was used.

### Whole genome functional screen using transposon directed insertion site sequencing (TraDIS)

The transposon mutant library was challenged with six bacteriophages (SPLA1a, SPLA1b, SPLA2, SPLA5B, SPLA5c and SPLA11) that exhibited full lytic activity against the wild type strain *S.* Typhimurium ST4/74. The Tn5 insertion mutant library was cultured in LB broth at 37°C with shaking for 18 hours. This culture was diluted to 1 x 10 ^7^ CFU/ml in 10ml of LB broth containing phage added at an MOI of 10, except SPLA1a which was added an MOI of 1, and incubated at 37 ⁰C for 3 hours. A buffer only negative control containing no phage supplementation was included. A 2 ml sample of each culture was then harvested by centrifugation at 2500 x g and genomic DNA extracted as previously described. Two independent replicates were performed.

### Preparation of DNA fragments for TraDIS sequencing

Genomic DNA from the transposon mutant library with and without exposure to SPLA phages was diluted to 11.1 ng/µl and tagmented using MuSeek DNA fragment library preparation kit (ThermoFisher, USA).

Fragmented DNA was purified using AMPure XP (Beckman Coulter, USA). DNA was amplified by PCR using biotinylated primers specific to the transposon and primers for the tagmented ends of DNA. PCR products were purified again using AMPure XP beads and incubated for 4 hours with streptavidin beads (Dynabeads®) to allow for capture of the DNA fragments with the transposon. A subsequent PCR step using barcoded sequencing primers allowed for the pooling of samples. Streptavidin beads were magnetically removed from the PCR products which were further purified and size-selected using AMPure XP beads. PCR products were quantified using Qubit 3.0 (Invitrogen, USA) and Tapestation (Aligent Technologies, USA).

The PCR products were then applied to a NextSeq 500 sequencing machine fitted with a NextSeq 500/550 High Output Kit v2.5 (75 Cycles) (Illumina). The nucleotide sequence reads obtained were then analysed using the BioTraDIS (30) software suite, which aligns the sequence reads to the *Salmonella enterica* sv Typhimurium strain ST4/74 reference genome nucleotide sequence, thereby identifying the location of transposon insertions and the number of reads that match at each site. This provides an approximate number of mutants at each site. Comparison with the reference genome annotation then provides mutant information for every gene, allowing fold changes (expressed as log _2_ fold change) and statistical significance (*q*-values) to be calculated between experimental conditions for each gene using. Data where *q* < 0.05 were considered to be statistically significant.

## Results

### Isolation of *Salmonella enterica* bacteriophages

To establish a diverse collection of phages capable of lysis of *S. enterica* strains, we enriched phages from wastewater, river or food samples using a diverse mixture of *S. enterica* enrichment strains in broth culture. Plaques with distinct morphologies were picked and purified further by rounds of single plaque purification on the preferred host strain. In total, 12 phages were isolated and designated SPLA1a, SPLA1b, SPLA2, SPLA3, SPLA4, SPLA5a, SPLA5b, SPLA5c, SPLA9, SPLA10, SPLA11 andSPLA12 (Table 1). Illumina short read sequencing of nucleic acid prepared from each phage lysate assembled into a single contiguous sequence for each phage, with contigs ranging in size from 40,585 bp to 240,593 bp. Except for SPLA1b, whole genome sequence reads of the *S. enterica* strains used to enrich the phage did not align to the assembled phage genomes indicating that these were not prophage derived from the enrichment strains. Sequence reads corresponding to a prophage in *S.* Typhimurium strain B8 C7 aligned to the full length of the SPLA1b assembled sequence indicating that this phage was likely activated from the enrichment strain.

**Table 1.** Summary of SPLA bacteriophage characteristics.

|  | Isolation Host<br>Serovar | Source | Genome Size<br>(bp) | Predicted Family | Predicted Genus |
| --- | --- | --- | --- | --- | --- |
| **SPLA1a** | S.Typhimurium | Food | 240348 | *Unclassified* | *Seoulvirus* |
| **SPLA1b** | S.Typhimurium | Food | 39459 | *Unclassified* | *Lederbergvirus* |
| **SPLA2** | S.Newport | Food | 53075 | *Unclassified* | *Rosemountvirus* |
| **SPLA3** | S.Mbandaka | River | 240593 | *Unclassified* | *Seoulvirus* |
| **SPLA4** | S.Mbandaka | Factory drain | 39540 | *Autographiviridae* | *Berlinvirus* |
| **SPLA5a** | S.Montevideo | Wastewater influent | 151208 | *Unclassified* | *Seunavirus* |
| **SPLA5b** | S.Kedougou | Wastewater influent | 107599 | *Demerecviridae* | *Tequintavirus* |
| **SPLA5c** | S.Infantis | Wastewater influent | 239099 | *Unclassified* | *Seoulvirus* |
| **SPLA9** | S.Montevideo | Wastewater influent | 51694 | *Unclassified* | *Rosemountvirus* |
| **SPLA10** | S.Montevideo | Wastewater influent | 210197 | *Unclassified* | *Phikzvirus* |
| **SPLA11** | S.Panama | Wastewater influent | 52451 | *Unclassified* | *Rosemountvirus* |
| **SPLA12** | S.Infantis | Wastewater influent | 51974 | *Unclassified* | *Rosemountvirus* |

### SPLA phages are diverse and represent seven distinct phage lineages

To investigate the diversity and relationship to known phage genome sequences in available databases SPLA phages were placed in a phylogenetic context based on sequence similarity of their predicted proteome sequence. Phylogenetic reconstruction using variation in the predicted proteome of SPLA phages with phage sequences in the Virus-host database indicated that all SPLA phages clustered with phages known to infect Gammaproteobacteria (Supplementary Figure 2). Therefore, to improve clarity, the analysis was repeated with only phages of Gammaproteobacteria (Figure 1). The SPLA phages were identified as members of the class *Caudoviricetes* and were present in clusters on seven deeply rooted lineages that corresponded to the genera *Berlinvirus*, *Seoulvirus*, *Phikzvirus*, *Tequintavirus*, *Seunavirus*, *Rosemountvirus* and *Lederbervirus*. SPLA1a, SPLA3 and SPLA5b were predicted to be myophages of the genus *Seoulvirus* with genome sizes of above 200kb and as such classified as jumbo phages (31). SPLA1a, SPLA3 and SPLA5b, of the genus *Seoulvirus*, had an average nucleotide sequence identity (ANI) greater than 98.4% and should be considered strains of the same species. SPLA2, SPLA9 and SPLA11 and SPLA12 were assigned to the genus *Rosemountvirus* and exhibited greater than 95.6% ANI. Phages SPLA4 and SPLA5b were the only phages which are currently assigned to established viral families, *Autographiviridae* and *Demerecviridae* respectively, according to the ICTV database (accessed December 2022).

**Figure 1.**
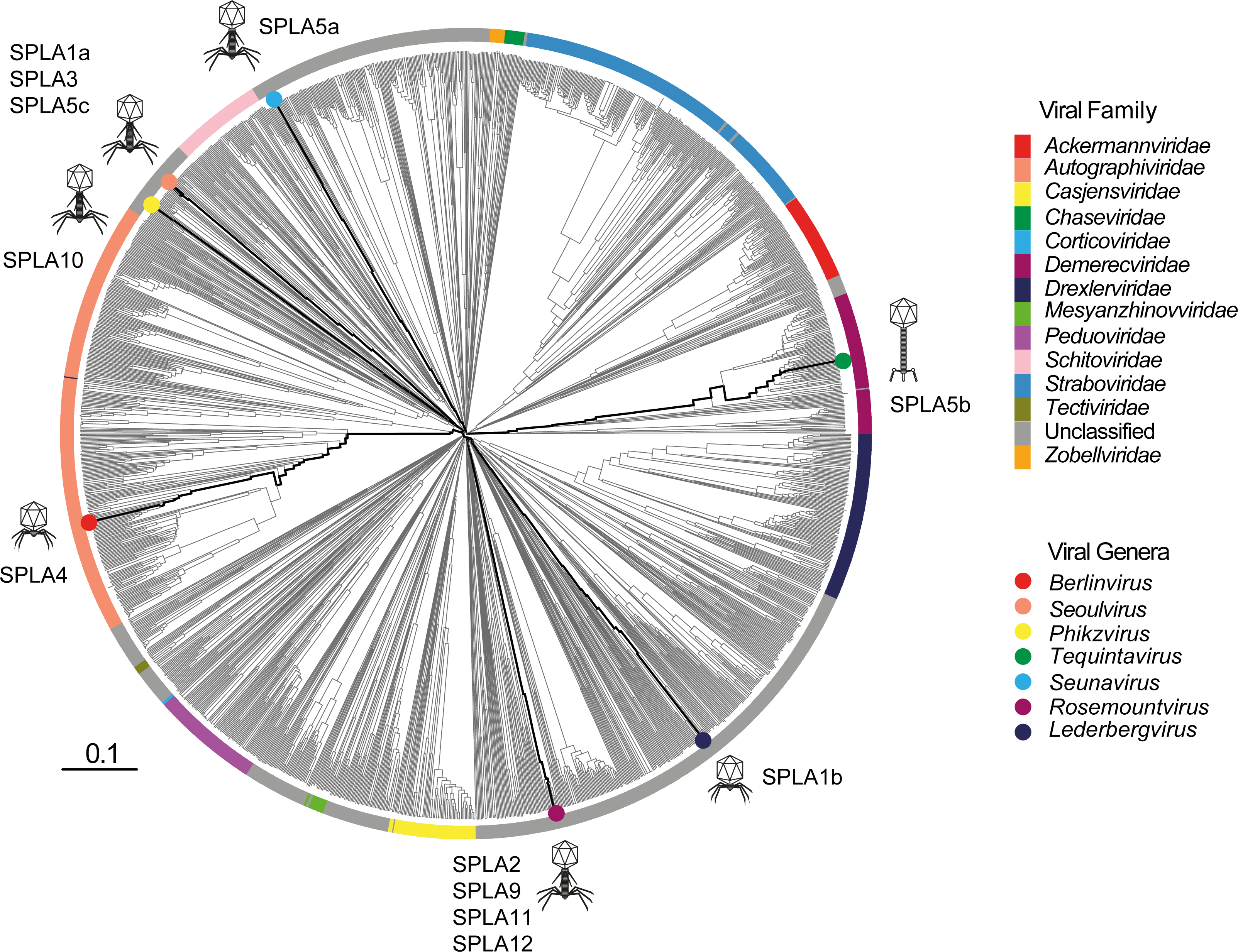
Relationship of SPLA phages proteome in the context of known diversity of viral proteomes from viral families known to infect gammaproteobacteria. . The dendrogram was generated using the proteome data with VipTree software and Viral Family was predicted using Inphared. SPLA *Salmonella* Phage Isolates with circles coloured by the viral genera Berlinvirus (red), Seoulvirus (orange), Phikzvirus (yellow), Tequintavirus (green), Seunavirus (Lightblue), Rosemountvirus (Purple) and Lederbergvirus (darkblue). Icons indicate predicted phage morphology.

### SPLA phages are highly specific to *Salmonella* serovars

To determine the host range of the SPLA phages, susceptibility of 36 bacterial strains of eleven diverse Gammaproteobacterial species were tested (Figure 2a). These included a representative strain from each of ten species of Enterobacterales, a strain of <u>Aeromonaceae</u> and 25 strains *S. enterica* comprising fourteen different serovars (Figure 2a). Strains of species other than *S. enterica* exhibited little susceptibility to SPLA phages indicated by the liquid assay score (LAS, 0-10) except for moderate susceptibility of a *Hafnia alvei*strain to several SPLA phages, particularly to SPLA2 and SPLA4 (Figure 2b). Susceptibility of *S. enterica* strains to SPLA phages varied markedly, with all phages exhibiting moderate (33-66 LAS) to high virulence (>66 LAS) to one or more strains of all 14 serovars tested. The taxonomically diverse phages SPLA9, SPLA11 and SPLA1a, exhibited the broadest host range, displaying at least moderate virulence for greater than nine of the fourteen serovars tested (Figure 2b and c). Nonetheless, all except SPLA5a, SPLA5b and SPLA1b had at least moderate virulence against at least six of the serovars tested (Figure 2b and c). Despite the close relationship of *S*. Typhimurium strains relative to strains from distinct serovars, SPLA phages exhibited similar variability in virulence. Phage SPLA1a was particularly virulent for a broad range of *S.* T y p h i m u r i u m strains and SPLA9 and SPLA11 were also moderately virulent for at least eleven of the 12 strains tested. Notably, even very closely related *Salmonella*strains exhibited diversity in sensitivity to SPLA phages. For example, *S*. Typhimurium strains S04698-09 and A53 were both part of the monophasic *S*. Typhimurium ST34 epidemic clade that emerged in the last three decades (32, 33), yet these strains exhibited distinct differences in susceptibility to at least five SPLA phages. Furthermore, a *S*. Typhimurium strain of ST4/74 that was genetically modified to remove prophage elements from its genome exhibited a moderate decrease in sensitivity to several SPLA phages.

**Figure 2.**
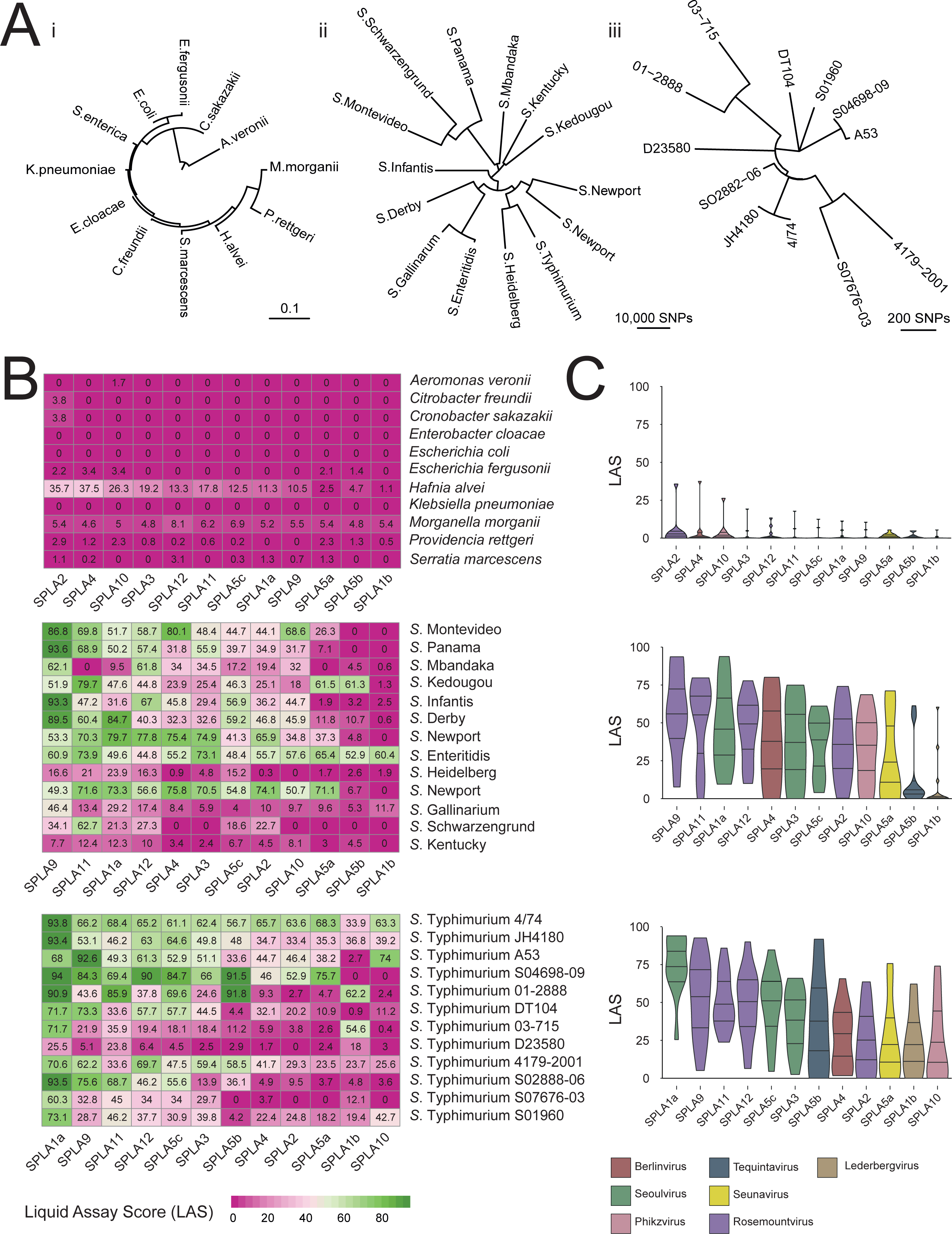
Host Range of SPLA phages for strains of diverse Gammaproteobacterial species and strains of *S. enterica*. A) Maximum likelihood phylogenetic trees showing the relationship and genetic distance of bacterial strains used for determination of host range based on single nucleotide polymorphisms (SNPs) in the 16S rRNA sequence of twelve strains of Proteobacterial species (i) and core genome of thirteen serovars (ii) and twelve strains of serovar Typhimurium (iii). B) Heat map showing liquid assay score (LAS) ranging from 0 (no change in growth) to 100 (no growth) of each phage-host bacterial strain combination of twelve strains of Proteobacterial species (top) and core genome of thirteen serovars (centre) and twelve strains of serovar Typhimurium (bottom). C) Violin plots of LAS scores for twelve strains of Proteobacterial species (top) and core genome of thirteen serovars (centre) and twelve strains of serovar Typhimurium (bottom) in the presence of each SPLA phage.

### Identification of SPLA phage host susceptibility and resistance genes using a functional genomics screen

To identify bacterial genes affecting the virulence of infecting SPLA phages, we constructed a Tn5 transposon insertion mutant library in *S*. Typhimurium strain ST4/74 comprising over 600,000 unique insertion sites corresponding, on average, at least one insertion every seven base pairs of the genome sequence. In a preliminary screen to determine susceptibility of the wild type strain of the transposon mutant library to the SPLA phages, we found that SPLA1a, SPLA1b, SPLA2, SPLA5b, SPLA5c and SPLA11 had a LAS > 50, indicating moderate to high virulence of the phages in question.

TraDIS was used to identify mutants whose abundance changed following cultures with each of these six bacteriophages compared to a control without phage treatment. Genes for which insertional inactivation mutants increased in frequency showed reduced phage susceptibility were termed susceptibility genes since their inactivation improved bacterial survival. Most of the susceptibility genes identified were genes which encoded proteins or were involved in the biosynthesis of macromolecules present on the outer surface of the bacterium (Figure 3). For example, insertional inactivation of *btuB,* that encodes a vitamin B12 (cobalamin) transporter was identified as a susceptibility gene for infection by phage SPLA5b (Figure 3D), while genes such as those in the *rfa* and *rfb* loci involved in the biosynthesis of lipopolysaccharide (LPS) were identified for SPLA1a, SPLA1b and SPLA5c (Figures 3A, 3B and E). Likewise, inactivation of genes including *yhjU*, *yhjL*, *yhjN*, *yhjQ*, *yhjR* and *yhjS,* involved in the biosynthesis of cellulose, reduced susceptibility to SPLA2 (Figure 3C).

**Figure 3.**
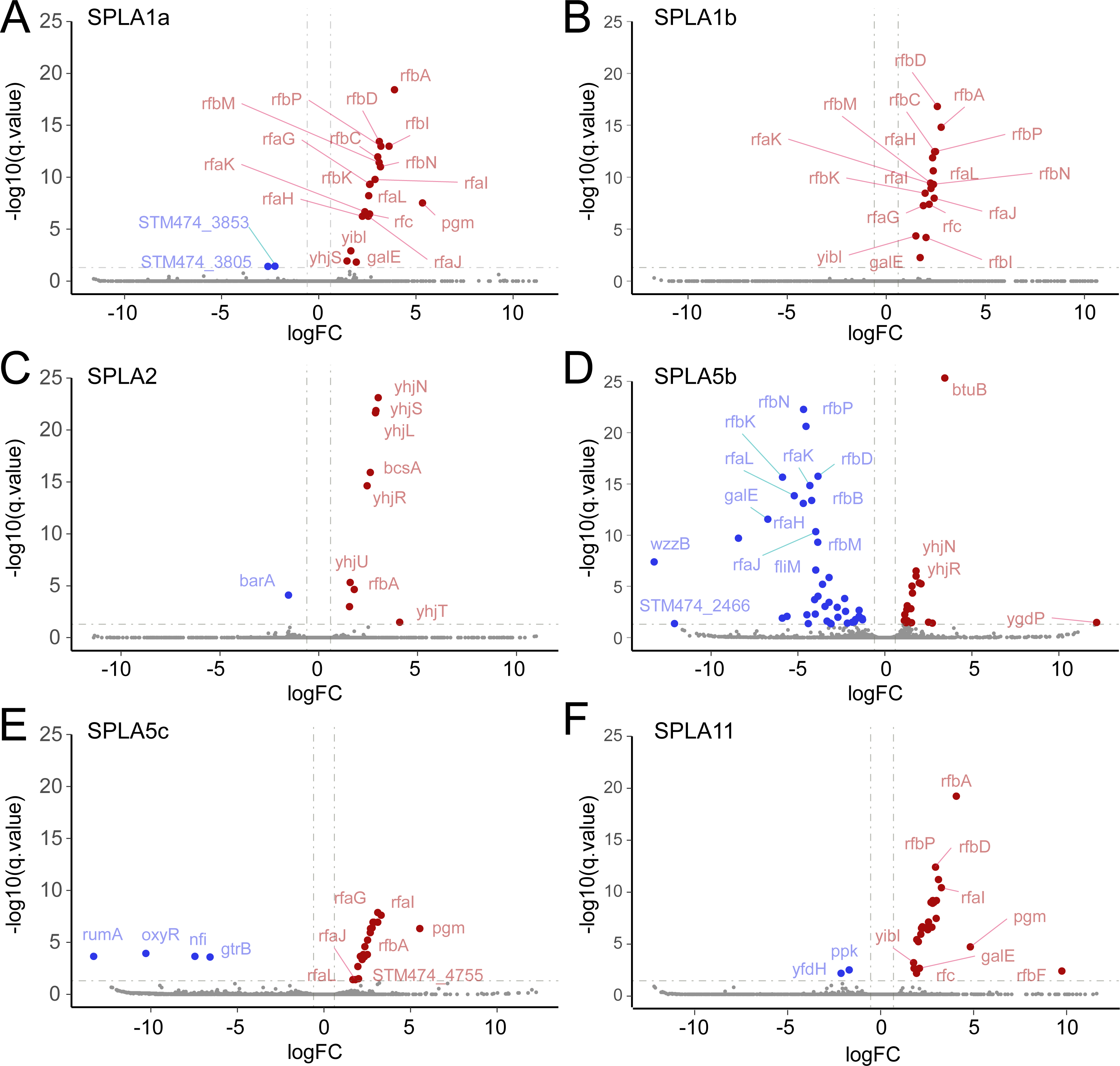
Changes in the Transposon Insertions within genes in response to predation by six SPLA phages. Log2 fold-change in insertions (x-axis) is plotted against -log _10_(q-value) statistical significance (y-axis) of difference in the number of insertions in each gene in cultures of S. Typhimurium treated with phage or untreated. Each graph shows gene hits for treatment with phage SPLA1a (A), SPLA1b (B), SPLA2 (C), SPLA5b (D), SPLA5c (E) and SPLA11 (F). Genes with significantly fewer insertions (blue points) or greater insertions (red points) are indicated. Selected genes are labelled with lines indicating the relevant data point.

In contrast, genes in which insertional inactivation mutants decreased in frequency following selection by SPLA phages were determined to provide resistance to bacteriophages. Putative resistance genes included two regulatory genes *barA* (SPLA2) and *oxyR* (SPLA5 c*r*)*u*,*mA* involved in methylation of uracil in 23S ribosomal RNA (SPLA5c), *nfi* encoding an endonuclease (SPLA5c) and *gtrB* involved in glycosylation of LPS (SPLA5c and SPLA11). Notably, while genes involved in synthesis of LPS were susceptibility genes for SPLA1a, SPLA1b and SPLA5c, these genes were resistance genes for infection by SPLA5b.

Notably, while insertional inactivation of genes coding for synthesis of LPS increased susceptibility to phages SPLA1a, SPLA1b and SPLA5c, it reduced susceptibility to phage SPLA5b.

### Closely related seoulviruses SPLA1a and SPLA5c differ in virulence due to sensitivity to O-antigen glycosylation by GtrB

SPLA1a and SPLA5c exhibited distinct virulence from each other to a range of strains of various *S. enterica* serovars, and to a range of strains of serovar *S*. Typhimurium (Figure 2), despite sharing 97% nucleotide sequence identity across their whole genome. For example, SPLA1a exhibited high virulence (LAS = 94) while SPLA5c exhibited moderate virulence (LAS = 61) for *S*. Typhimurium strain ST4/74. A key functional genomic difference between SPLA1a and SPLA5c was that virulence of SPLA5c was sensitive to the presence of t h*g*e*trB* gene, encoding a glycosyltransferase. A *S*. Typhimurium strain ST4/74 in which the *gtrB* gene was replaced by an *aphII* gene resulted in an increase in LAS to 100 for SPLA5c, indicating virulence comparable to that for SPLA1a (Figure 4). To investigate differences in nucleotide sequence of SPLA1a and SPLA5c that may account for the distinct sensitivity of each phage to the *gtrB* phage resistance gene, the genomes were aligned and regional variation in nucleotide sequence identity and insertions or deletion were identified. SPLA1a had two insertions relative to SPLA5c affecting a gene of unknown function and a region encoding a putative virion structural protein. SPLA5c had three insertions affecting a gene encoding a putative RNA polymerase subunit, the terminase large subunit, and a protein of unknown function (Supplementary Figure 3). There was also a large region of approximately 10 kb with greater sequence divergence affecting 17 genes that mostly encoded proteins of unknown function, but also virion structural proteins and a putative tail fibre protein.

**Figure 4.**
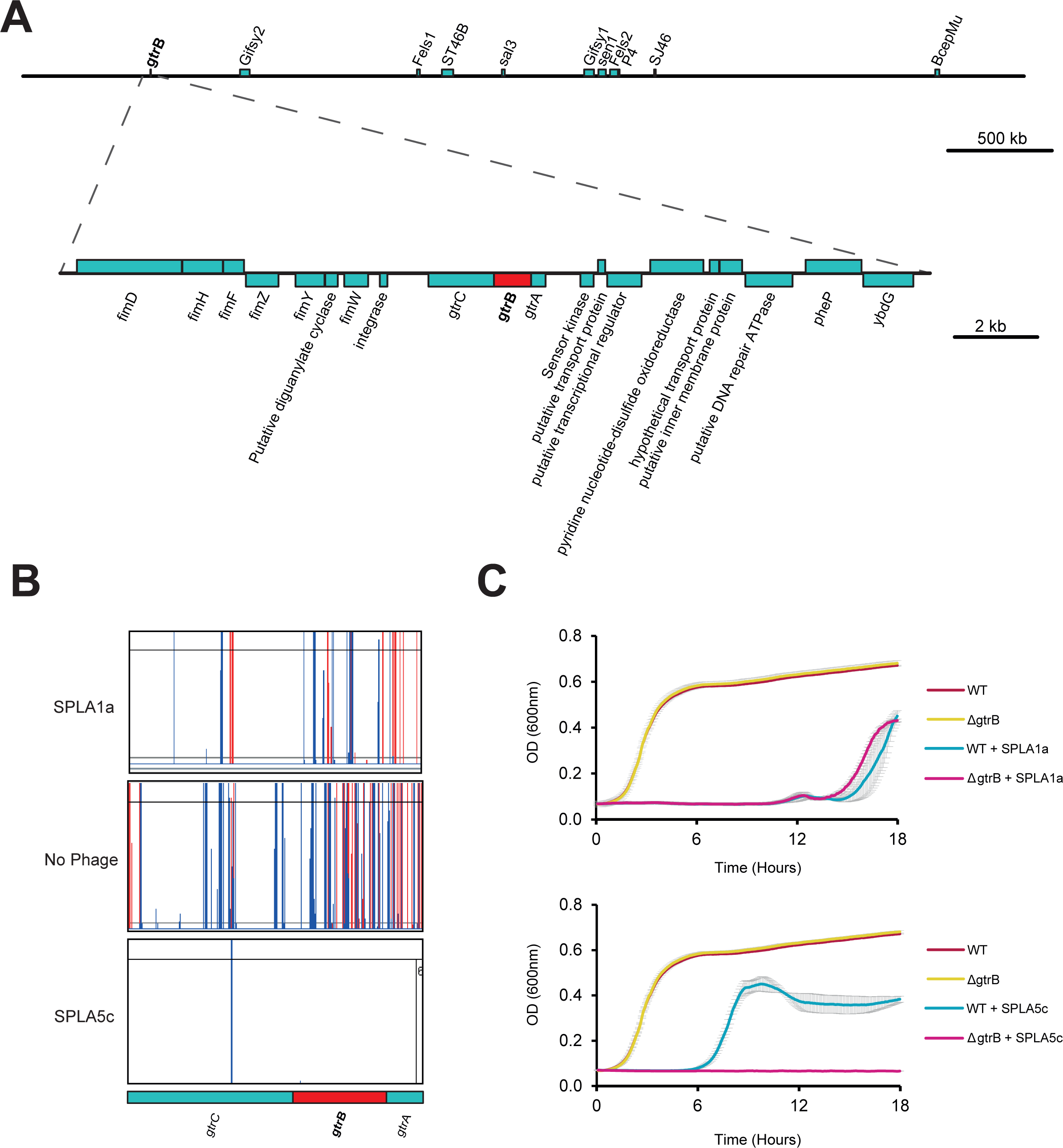
The role of *gtrB* in phage resistance to SPLA5c. Genomic context of *gtrABC* genes in *Salmonella* Typhimurium ST4/74 with gtrB (red bar) and neighbouring genes (teal) indicated on forward and reverse strands above an below horizontal line (A). Transposon Insertion Sites mapped to *Salmonella* Typhimurium 4/74 in the *gtrABC* operon in the presence of SPLA1a, SPLA5c and non-Phage Treated control. The height of the vertical lines indicates the number of insertions of Tn5 detected in the forward (red) and reverse (blue) orientation in the *gtrABC* genes are shown. Growth of *S*. Typhimurium strain 4/74 and an otherwise isogenic strain in which the *gtrB* gene has been replaced by the *aphII* gene (Δ*gtrB*) in the presence or absence of SPLA1a (top graph) and SPLA5c (bottom graph).

### Functional genomics predicts collateral sensitivity of resistant escape mutants and enables rational design of phage combinations with increased efficacy

Collateral sensitivity is an evolutionary trade-off where resistance to one antimicrobial agent results in increased sensitivity to another (34). We observed that LPS biosynthesis genes played contrasting roles as resistance genes or sensitivity genes during infection with phage SPLA1a and SPLA5b, respectively. To investigate the differential role of LPS in virulence of SPLA1a and SPLA5b phages their virulence was determined for otherwise isogenic *S*. Typhimurium ST4/74 strains in which *rfaL* or *btuB* were deleted. Both SPLA1a and SPLA5b were able to form plaques on wild-type *Salmonella* Typhimurium ST4/74, although plaques formed by SPLA5b were more turbid than those formed by SPLA1a, consistent with a lower virulence of the former (Figure 5a). Deletion of *rfaL* resulted in complete resistance to SPLA1a, as indicated by the lack of plaques in overlay assays, consistent with O-antigen being the receptor for this phage. In contrast, SPLA5b was still able to infect the *rfaL* mutant, as expected from the TraDIS data, but plaques were markedly less turbid, suggesting an increase in phage sensitivity. In contrast, loss of the vitamin B12 transporter, by disruption of the *btuB* gene, resulted in resistance to SPLA5b as indicated by the lack of plaques, but no effect on SPLA1a infection (Figure 5). This was consistent with BtuB being the receptor for SPLA5b.

**Figure 5.**
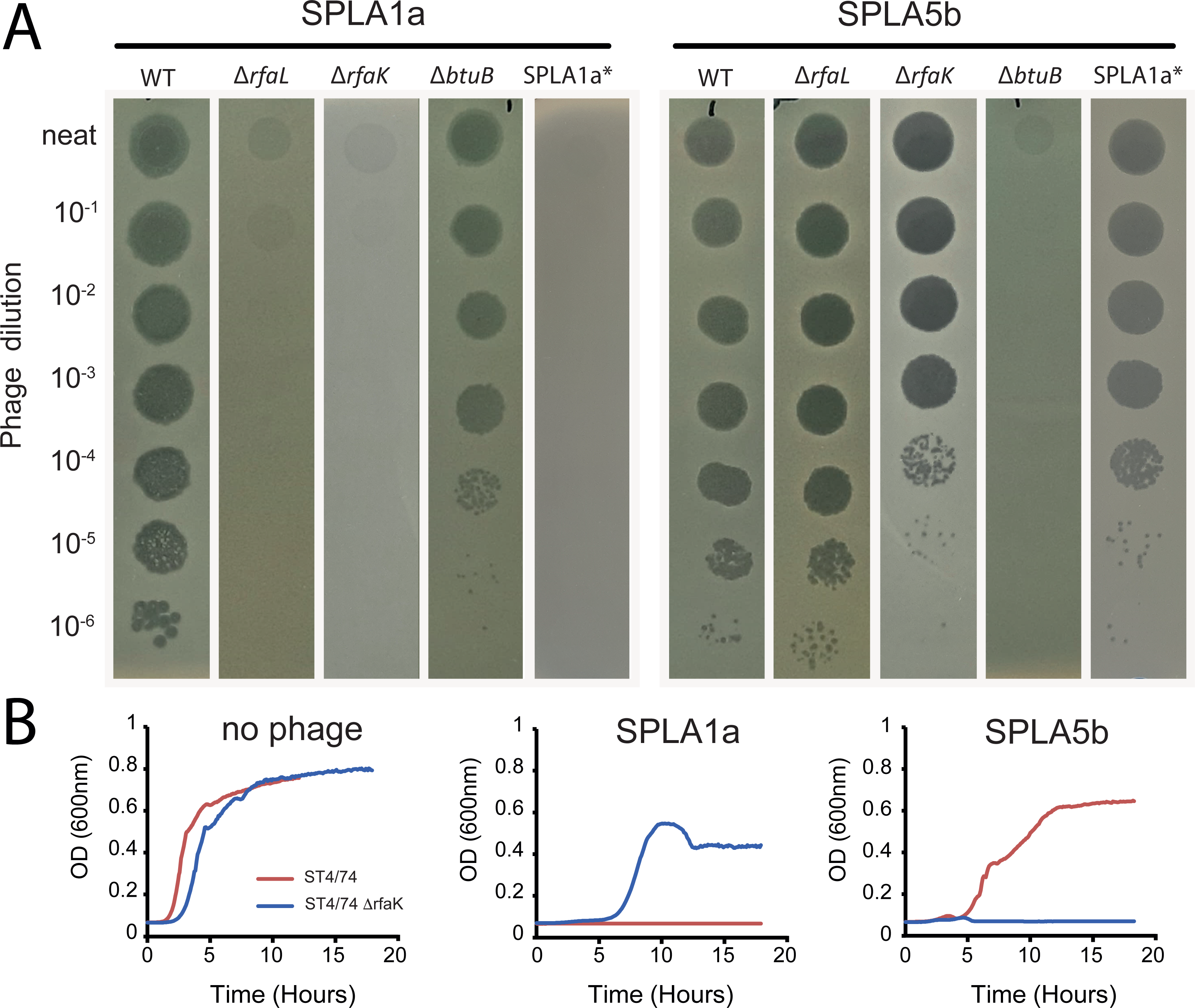
Inactivation of genes in S. Typhimurium strain ST4/74 that confer resistance to SPLA1a result in increased susceptibility to SPLA5b phage. Collateral sensitivity dynamic of Bacteriophages SPLA1a and SPLA5b. Plaque assays using SPLA1a (left) and SPLA5b (Right) against LPS and BtuB Mutants (A) and growth in broth culture of S. Typhimurium strain ST4/74 (Red) or an otherwise isogenic strain in which the *rfaK* genes was replaced by tetA gene (ΔrfaK, (blue) when treated with no phages (left graph), SPLA1a (middle graph) and SPLA5b (right graph) (B).

To further determine if loss of LPS resulted in increased susceptibility to SPLA5b, the virulence of SPLA phages on ST4/74 strain lacking the *rfaK* gene (Δ *rfaK*) that encodes a hexose transferase involved in LPS synthesis, was determined. No growth of wild-type ST4/74 was observed in the presence of SPLA1a, but strains with a deletion in *rfaK* resulted in the ability to grow under SPLA1a selection. In contrast, growth was observed when wild-type ST4/74 was infected with SPLA5b, with the phage causing an extended lag phase compared to non-phage treated conditions. However, loss of *rfaK* resulted in increased phage infection by SPLA5b, compared to the wild-type (Figure 5b).

Since mutations in the same receptor were confirmed to play contrasting roles in infection by SPLA1a and SPLA5b, it raised the possibility that naturally occurring resistance to SPLA1a may result in increased sensitivity to SPLA5b. Therefore, collateral sensitivity may improve the efficacy of these phages in combination treatment. In cultures of *S*. Typhimurium ST4/74 infected with SPLA1a, approximately 10% (10/96) resulted in the emergence of naturally occurring resistance within 18 hours of incubation at 37 °C (Figure 6). Whole genome sequence of a resistant strain (4/74 SPLA1a*) revealed a single base insertion in the *pgm* gene that resulted in a frame shift predicted to truncate this gene (Supplementary Figure 4). The *pgm* gene, encoding phosphoglucomutase, was also identified as a host susceptibility gene for SPLA1a phage in the transposon mutant library screen (Figure 3A). In contrast to SPLA1a, upon infection of *S*. Typhimurium ST4/74 with SPLA5b host resistance emerged in 100% (96/96) of cultures, although growth was delayed, and the final optical density was lower than uninfected controls (Figure 6). Consistent with the idea that resistance to SPLA1a results in collateral sensitivity to SPLA5b, both phages were used in combination and growth was only observed in 1/96 of cultures, an approximate 10-fold reduction in phage resistance compared to SPLA1a alone. This reduction was consistently observed across five biological replicates.

**Figure 6.**
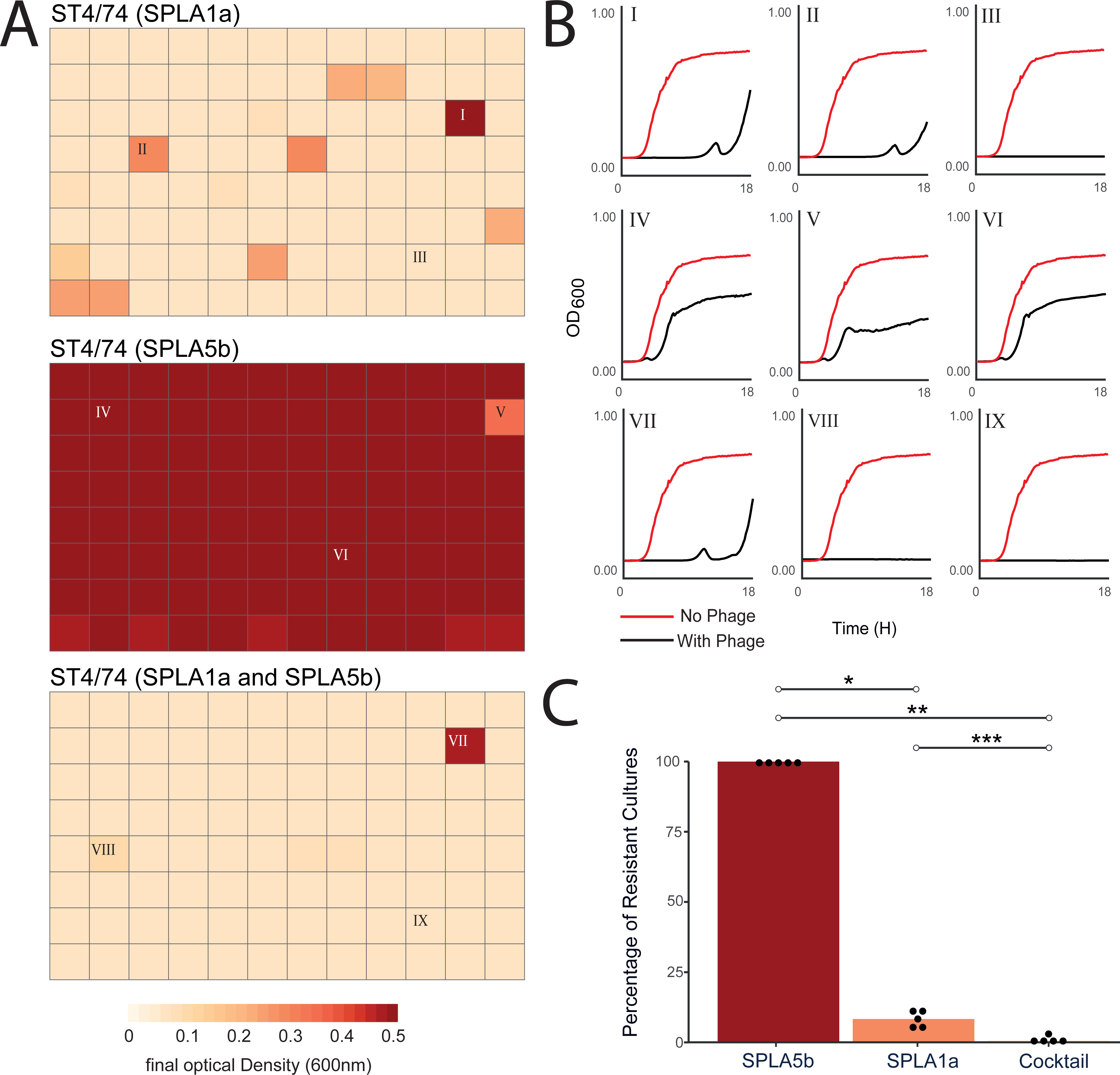
Growth of *Salmonella* Typhimurium ST4/74 in the presence of SPLA1a and SPLA5b in monotherapy and combination therapy. The end point optical density at 600nm (OD_600nm_) after 18 hours culture of S. Typhimurium strain ST4/74 in the presence of phage SPLA1a (top), phage SPLA5b (middle) or SPLA1a and SPLA5b (bottom) (A). Example growth curves of *Salmonella* Typhimurium ST4/74 infected by SPLA1a (I-III) SPLA5b (IV-VI) and SPLA1a and SPLA5b (VII-IX) for which roman numerals correspond to culture end points labelled previously (B). Percentage of resistant cultures for SPLA1a, SPLA5b and combination treatment across 5 biological replicates. Statistical significance is shown with (*) (C)

We propose that resistance to SPLA1a due to mutations affecting production of LPS results in a collateral sensitivity dynamic, increasing sensitivity to SPLA5b. A working hypothesis that is consistent with our observations is that wild-type*S*. Typhimurium ST4/74 strains that elaborate LPS with long chain O-antigen on their surface that masks the BtuB receptor used by SPLA5b for initiation of infection. Mutations in LPS biosynthesis, such as Δ *rfaL*, lead to resistance to SPLA1a provide greater access to the BtuB receptor increased sensitivity to SPLA5b (Figure 7).

**Figure 7.**
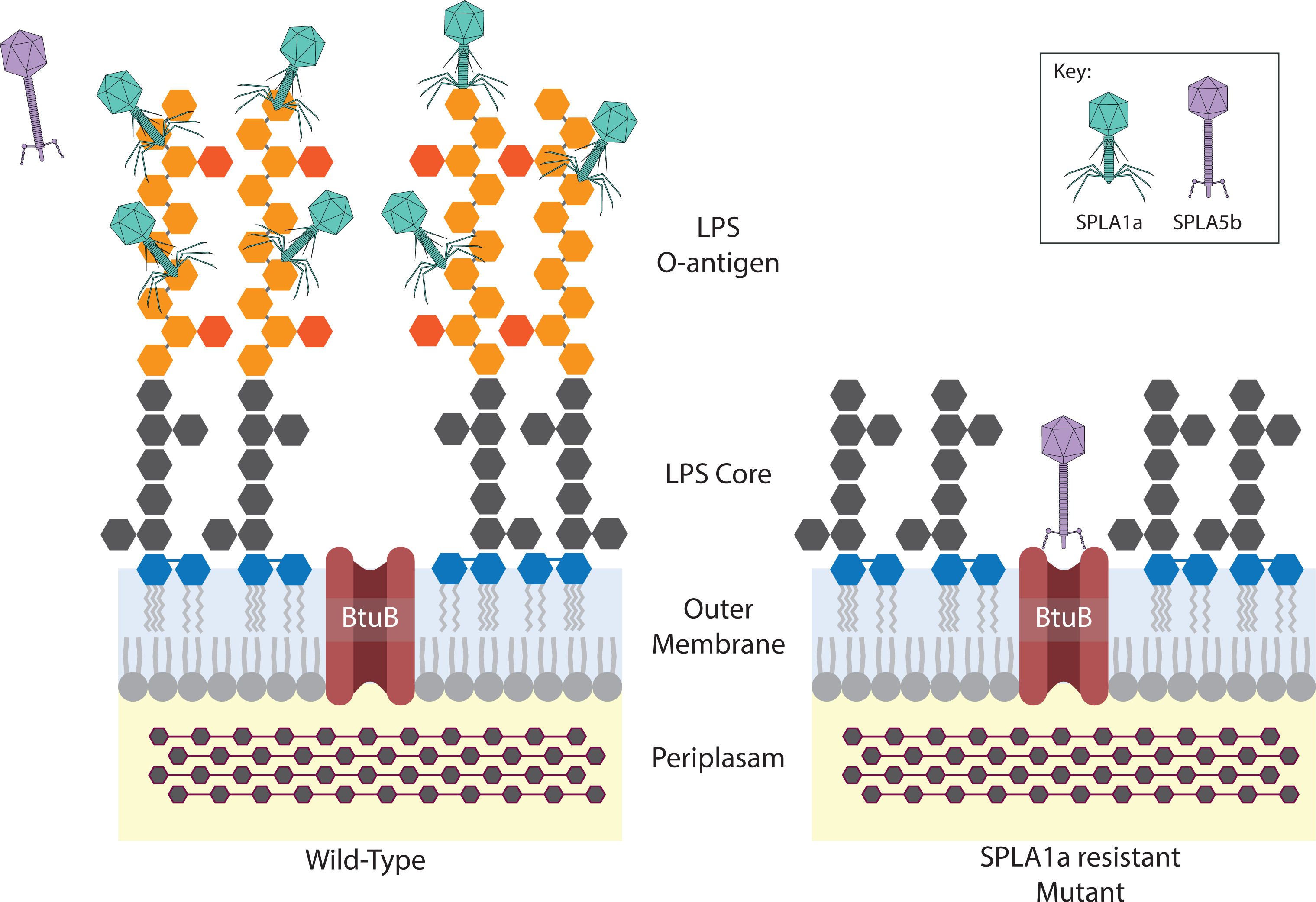
Schematic of predicted collateral sensitivity mechanism observed in combination treatment with SPLA1a and SPLA5b. Wild-type strains (left) are highly susceptible to SPLA1a, but less susceptible to SPLA5b due to receptor masking. SPLA1a insensitive mutant (right) is no longer infected by SPLA1a, due to lack of O-antigen receptor, but has increased susceptibility to SPLA5b infection due to loss of receptor masking.

## Discussion

Phages show potential as antimicrobials to control bacterial pathogens in the environment or to replace or augment therapies to treat infections. Two of the main technological challenges to the development of phage-based products is their specificity and the rapid emergence of resistance by target bacteria. Although strain or species specificity is a potential advantage, as bystander beneficial bacteria are not affected during application of the bacteriophage, specificity also presents a challenge to the development of broadly applicable products. A common solution is to formulate a cocktail of several phages with diverse host ranges and receptors. This approach may also contribute to decreasing the likelihood of the emergence of resistance since multiple mutations may be required to produce resistance to the cocktail of phage. Previous studies have shown that phage cocktails are effective against many pathogens including *Enterococcus*, *Salmonella* a n d *Campylobacter* (35–37). In this study we devised a functional genomics approach to direct the formulation of combinations of newly isolated phages, using a transposon mutant screen in conjunction with TraDIS to identify genes that either increased sensitivity, susceptibility genes, and genes that decreased sensitivity to genes, resistance genes.

The formulation of optimal phage cocktails is likely to depend on combining phylogenetically diverse phage, as related phages are likely to use identical or similar bacterial encoded receptors to initiate infection. Furthermore, their susceptibility to resistance mechanisms encoded by the target bacteria is likely to be shared. We isolated 12 phage that represented seven different viral genera, based on whole genome sequence and proteome comparison. All phages had limited virulence for strains of species other than *Salmonella enterica*, signifying their potential suitability for applications where specificity is important. Notably, the greatest off-target virulence was observed for a strain of *Hafnia alviei*, a species that is relatively distantly related to *Salmonella enterica*, while all phage were completely avirulent for *E. coli* that is relatively closely related to *Salmonella enterica*.

Considerable variation in virulence of all phages was observed for representative strains of 14 serovars of *Salmonella enterica*, highlighting the importance of combinations of phage to ensure broad activity. One or more phages showed moderate to high virulence to serovars Typhimurium, Newport, Derby that together account for 89% of pig isolates in 2020 and serovars Kedougou, Montevideo, Mbandaka, Newport and Enteritidis that together accounted for 46% of poultry isolates in 2020 (38), supporting their potential for development as antimicrobials in the food chain. A similar level of variation in virulence of SPLA phages for eleven strains of *S*. Typhimurium to that for representative strains of diverse serovars was observed. This was perhaps surprising since *S*. Typhimurium exhibits less than 0.1% nucleotide sequence divergence in their genome (39), compared with up to 2% nucleotide sequence divergence for strains of *S. enterica* (32). *S*. Typhimurium accounts for almost one quarter of all *Salmonella* isolated from humans and 78% of pig isolates in 2020 in the UK (38), this serovar is of particular interest for interventions. *S.* Typhimurium is particularly widespread in nature, being in the top three serovars isolated from multiple species of livestock and wild animals and is also relatively genetically diverse compared to other serovars (40) with multiple lineages exhibiting broad host range or host adaptation (41). This level of diversity has not been observed for other serovars and it remains to be determined if a similar level of variation in sensitivity to phage also occurs in strains of serovars other than Typhimurium.

Phage SPLA1a was particularly virulent for multiple *S*. Typhimurium strains, exhibiting high virulence for all but strain D23580. Strain D23580 is from a clade of sequence type (ST) 313 associated with invasive disease in sub-Saharan Africa and is rarely associated with infection outside of this geographical location (42, 43). Other SPLA phages also exhibited low virulence or were unable to infect strain D23580. This strain has a unique prophage repertoire including BTP-1 and BTP-5 (44, 45). BTP-1 harbours the phage resistance gene *bstA* which confers population-level resistance to bacteriophages via abortive infection (46) and may explain the relative phage resistance observed in this strain. SPLA1a was also the third most virulent SPLA phage for strains from diverse serovars of *S. enterica* and is therefore a potential key component of phage cocktails. Despite the relatively high virulence of SPLA1a, approximately 10% of *S*. Typhimurium ST4/74 cultures infected with the phage exhibited some growth around 18 hours later.

Infection by bacteriophages is a complex process with host bacteria encoding many genes affecting infection (47). Transposon insertion mutant library screens have been used to identify host genes involved in infection by model phages such as for T4 (48), λ (49) T2, T6 and T7 Phages (50). Additionally, it has been used to identify capsule polysaccharides as the receptor for novel phage, RAD2, in *Klebsiella* (*51*) and Vi and genetic regulators of a *S*. Typhi phage (52). However, it is not common practice to identify host resistance and susceptibility genes during the development of phage cocktails (53). Three of five lytic phages investigated here, SPLA1a, SPLA5c and SPLA11 likely recognise the LPS as the primary receptor, while SPLA2 and SPLA5b recognise cellulose and BtuB, respectively. These susceptibility genes are candidate genes that may be inactivated in strains that become resistant on exposure to phage. A common mechanism of defence against O-antigen targeting phages is the deletion of LPS biosynthesis genes or epigenetic control of O-antigen chain length (54, 55). We also observed a mutation in the *pgm* gene within a resistant strain that emerged in a culture *S*. Typhimurium exposed to SPLA1a phage. No previously described specific resistance mechanisms were identified. However, genes encoding regulators OxyR and BarA that may repress expression of a phage receptor in the testculture condition were identified as candidate resistance genes for SPLA5c and SPLA2 respectively. Also, the *nfi* gene that is predicted to encode an endonuclease involved in DNA repair (56) may play a role in phage resistance to SPLA5c and genes that modified the O- antigen receptor by glycosylation thereby preventing access to the O-antigen receptor were identified. These genes are of potential interest as molecular markers of resistance that could be used to predict the outcome of phage therapy.

Of particular interest for rational design of phage cocktails was the observation that some genes were susceptibility genes for one group of phage and a resistance gene for a second phage. This was the case of LPS biosynthesis genes that were susceptibility genes for SPLA1a, SPLA5c and SPLA11, and resistance genes for SPLA5b. Mutations in susceptibility genes identified in our transposon insertion mutant library are candidate genes that may be inactivated in strains that become resistant on exposure to phage. In order to exploit this resistance mutation, we tested the efficacy of a mixture of SPLA1a with SPLA5b and found that addition of SPLA5b increased efficacy by 10-fold.

Perhaps the greatest challenge for successful implementation of phage therapy are concerns of the rapid acquisition of phage resistance and treatment failure. Our data support the idea that the use of functional genomic screens is effective in identifying phage that target phage resistant escape mutants of additional phage facilitating the design of cocktails with improved efficacy. Additionally, identification of the genetic determinants of phage susceptibility can be used to help predict which strains a phage or phage cocktail can infect, which is likely to be important for successful implementation of phage therapy.

## References

1. Thanki AM, Mignard G, Atterbury RJ, Barrow P, Millard AD, Clokie MR. 2022. Prophylactic delivery of a bacteriophage cocktail in feed significantly reduces Salmonella colonization in pigs. Microbiology spectrum 10:e00422–22.

2. Thanki AM, Hooton S, Gigante AM, Atterbury RJ, Clokie MR. 2021. Potential roles for bacteriophages in reducing Salmonella from poultry and swine, Salmonella spp-a global challenge. IntechOpen.

3. Pollock J, Low AS, McHugh RE, Muwonge A, Stevens MP, Corbishley A, Gally DL. 2020. Alternatives to antibiotics in a One Health context and the role genomics can play in reducing antimicrobial use. Clin Microbiol Infect 26:1617–1621.

4. Bertozzi Silva J, Storms Z, Sauvageau D. 2016. Host receptors for bacteriophage adsorption. FEMS Microbiology Letters 363.

5. Lood C, Haas P-J, van Noort V, Lavigne R. 2022. Shopping for phages? Unpacking design rules for therapeutic phage cocktails. Current Opinion in Virology 52:236–243.

6. Sarker SA, Sultana S, Reuteler G, Moine D, Descombes P, Charton F, Bourdin G, McCallin S, Ngom-Bru C, Neville T. 2016. Oral phage therapy of acute bacterial diarrhea with two coliphage preparations: a randomized trial in children from Bangladesh. EBioMedicine 4:124–137.

7. Thanki AM, Clavijo V, Healy K, Wilkinson RC, Sicheritz-Ponten T, Millard AD, Clokie MRJ. 2022. Development of a Phage Cocktail to Target Salmonella Strains Associated with Swine. Pharmaceuticals (Basel) 15.

8. Eng S-K, Pusparajah P, Ab Mutalib N-S, Ser H-L, Chan K-G, Lee L-H. 2015. Salmonella: a review on pathogenesis, epidemiology and antibiotic resistance. Frontiers in Life Science 8:284–293.

9. EFSA. 2014. The European Union Summary Report on Trends and Sources of Zoonoses, Zoonotic Agents and Food-borne Outbreaks in 2012. EFSA Journal 12:312.

10. Rodwell EV, Wenner N, Pulford CV, Cai Y, Bowers-Barnard A, Beckett A, Rigby J, Picton DM, Blower TR, Feasey NA. 2021. Isolation and characterisation of bacteriophages with activity against invasive non-typhoidal Salmonella causing bloodstream infection in Malawi. Viruses 13:478.

11. Datsenko KA, Wanner BL. 2000. One-step inactivation of chromosomal genes in Escherichia coli K-12 using PCR products. Proceedings of the National Academy of Sciences 97:6640–6645.

12. Datta S, Costantino N, Court DL. 2006. A set of recombineering plasmids for gram-negative bacteria. Gene 379:109–115.

13. Chan W, Costantino N, Li R, Lee SC, Su Q, Melvin D, Court DL, Liu P. 2007. A recombineering based approach for high-throughput conditional knockout targeting vector construction. Nucleic acids research 35:e64–e64.

14. Thomson NM, Zhang C, Trampari E, Pallen MJ. 2020. Creation of Golden Gate constructs for gene doctoring. BMC Biotechnol 20:54.

15. Clokie MR, Kropinski A. 2009. Methods and Protocols, Volume 1: Isolation, Characterization, and Interactions. Springer.

16. Chen S, Zhou Y, Chen Y, Gu J. 2018. fastp: an ultra-fast all-in-one FASTQ preprocessor. bioRxiv doi:10.1101/274100:274100.

17. Seeman T. 2018. Shovill: faster SPAdes assembly of Illumina reads (v0. 9.0).

18. Bankevich A, Nurk S, Antipov D, Gurevich AA, Dvorkin M, Kulikov AS, Lesin VM, Nikolenko SI, Pham S, Prjibelski AD. 2012. SPAdes: a new genome assembly algorithm and its applications to single-cell sequencing. Journal of computational biology 19:455–477.

19. Li H, Durbin R. 2010. Fast and accurate long-read alignment with Burrows–Wheeler transform. Bioinformatics 26:589–595.

20. Nishimura Y, Yoshida T, Kuronishi M, Uehara H, Ogata H, Goto S. 2017. ViPTree: the viral proteomic tree server. Bioinformatics 33:2379–2380.

21. Cook R, Brown N, Redgwell T, Rihtman B, Barnes M, Clokie M, Stekel DJ, Hobman J, Jones MA, Millard A. 2021. INfrastructure for a PHAge REference Database: Identification of Large-Scale Biases in the Current Collection of Cultured Phage Genomes. PHAGE 2:214–223.

22. Jain C, Rodriguez-R LM, Phillippy AM, Konstantinidis KT, Aluru S. 2018. High throughput ANI analysis of 90K prokaryotic genomes reveals clear species boundaries. Nature Communications 9:5114.

23. Katoh K, Misawa K, Kuma Ki, Miyata T. 2002. MAFFT: a novel method for rapid multiple sequence alignment based on fast Fourier transform. Nucleic acids research 30:3059–3066.

24. Capella-Gutiérrez S, Silla-Martínez JM, Gabaldón T. 2009. trimAl: a tool for automated alignment trimming in large-scale phylogenetic analyses. Bioinformatics 25:1972–1973.

25. Minh BQ, Schmidt HA, Chernomor O, Schrempf D, Woodhams MD, von Haeseler A, Lanfear R. 2020. IQ-TREE 2: New Models and Efficient Methods for Phylogenetic Inference in the Genomic Era. Molecular Biology and Evolution 37:1530–1534.

26. Kalyaanamoorthy S, Minh BQ, Wong TKF, von Haeseler A, Jermiin LS. 2017. ModelFinder: fast model selection for accurate phylogenetic estimates. Nature Methods 14:587–589.

27. Hoang DT, Chernomor O, von Haeseler A, Minh BQ, Vinh LS. 2018. UFBoot2: Improving the Ultrafast Bootstrap Approximation. Molecular Biology and Evolution 35:518–522.

28. Stamatakis A. 2014. RAxML version 8: a tool for phylogenetic analysis and post-analysis of large phylogenies. Bioinformatics 30:1312–1313.

29. Xie Y, Wahab L, Gill JJ. 2018. Development and Validation of a Microtiter Plate-Based Assay for Determination of Bacteriophage Host Range and Virulence. Viruses 10.

30. Barquist L, Mayho M, Cummins C, Cain AK, Boinett CJ, Page AJ, Langridge GC, Quail MA, Keane JA, Parkhill J .2016. The TraDIS toolkit: sequencing and analysis for dense transposon mutant libraries. Bioinformatics (Oxford, England) 32:1109–1111.

31. Yuan Y, Gao M. 2017. Jumbo Bacteriophages: An Overview. Frontiers in microbiology 8:403–403.

32. Pye HV, Thilliez G, Acton L, Kolenda R, Al-Khanaq H, Grove S, Kingsley RA. 2023. Strain and serovar variants of Salmonella enterica exhibit diverse tolerance to food chain-related stress. Food Microbiology 112:104237.

33. Petrovska L, Mather AE, AbuOun M, Branchu P, Harris SR, Connor T, Hopkins K, Underwood A, Lettini AA, Page A. 2016. Microevolution of monophasic Salmonella Typhimurium during epidemic, United Kingdom, 2005–2010. Emerging infectious diseases 22:617.

34. Pal C, Papp B, Lazar V. 2015. Collateral sensitivity of antibiotic-resistant microbes. Trends Microbiol 23:401–7.

35. Wandro S, Ghatbale P, Attai H, Hendrickson C, Samillano C, Suh J, Pride DT, Whiteson K. 2021. Phage Cocktails can Prevent the Evolution of Phage-Resistant Enterococcus. bioRxiv.

36. Fischer S, Kittler S, Klein G, Glünder G. 2013. Impact of a Single Phage and a Phage Cocktail Application in Broilers on Reduction of Campylobacter jejuni and Development of Resistance. PLOS ONE 8:e78543.

37. Pereira C, Moreirinha C, Lewicka M, Almeida P, Clemente C, Cunha Â, Delgadillo I, Romalde JL, Nunes ML, Almeida A. 2016. Bacteriophages with potential to inactivate Salmonella Typhimurium: Use of single phage suspensions and phage cocktails. Virus Research 220:179–192.

38. APHA. 2019. Salmonella In Livestock Production in GB, 2020. Animal and Plant Health Agency,

39. Branchu P, Bawn M, Kingsley RA. 2018. Genome variation and molecular epidemiology of Salmonella enterica serovar Typhimurium pathovariants. Infection and immunity 86:e00079–18.

40. Alikhan N-F, Zhou Z, Sergeant MJ, Achtman M. 2018. A genomic overview of the population structure of Salmonella. PLoS genetics 14:e1007261.

41. Bawn M, Alikhan N-F, Thilliez G, Kirkwood M, Wheeler NE, Petrovska L, Dallman TJ, Adriaenssens EM, Hall N, Kingsley RA. 2020. Evolution of Salmonella enterica serotype Typhimurium driven by anthropogenic selection and niche adaptation. PLoS genetics 16:e1008850.

42. Kingsley RA, Msefula CL, Thomson NR, Kariuki S, Holt KE, Gordon MA, Harris D, Clarke L, Whitehead S, Sangal V, Marsh K, Achtman M, Molyneux ME, Cormican M, Parkhill J, MacLennan CA, Heyderman RS, Dougan G. 2009. Epidemic multiple drug resistant Salmonella Typhimurium causing invasive disease in sub-Saharan Africa have a distinct genotype. Genome research 19:2279–2287.

43. Okoro CK, Kingsley RA, Connor TR, Harris SR, Parry CM, Al-Mashhadani MN, Kariuki S, Msefula CL, Gordon MA, de Pinna E, Wain J, Heyderman RS, Obaro S, Alonso PL, Mandomando I, MacLennan CA, Tapia MD, Levine MM, Tennant SM, Parkhill J, Dougan G. 2012. Intracontinental spread of human invasive Salmonella Typhimurium pathovariants in sub-Saharan Africa. Nat Genet 44:1215–21.

44. Owen SV, Wenner N, Canals R, Makumi A, Hammarlöf DL, Gordon MA, Aertsen A, Feasey NA, Hinton JCD. 2017. Characterization of the Prophage Repertoire of African Salmonella Typhimurium ST313 Reveals High Levels of Spontaneous Induction of Novel Phage BTP1. Frontiers in Microbiology 8.

45. Bawn M, Alikhan NF, Thilliez G, Kirkwood M, Wheeler NE, Petrovska L, Dallman TJ, Adriaenssens EM, Hall N, Kingsley RA. 2020. Evolution of Salmonella enterica serotype Typhimurium driven by anthropogenic selection and niche adaptation. PLoS Genet 16:e1008850.

46. Owen SV, Wenner N, Dulberger CL, Rodwell EV, Bowers-Barnard A, Quinones-Olvera N, Rigden DJ, Rubin EJ, Garner EC, Baym M, Hinton JCD. 2021. Prophages encode phage- defense systems with cognate self-immunity. Cell Host & Microbe 29:1620–1633.e8.

47. Azam AH, Tanji Y. 2019. Bacteriophage-host arm race: an update on the mechanism of phage resistance in bacteria and revenge of the phage with the perspective for phage therapy. Applied Microbiology and Biotechnology 103:2121–2131.

48. Cowley LA, Low AS, Pickard D, Boinett CJ, Dallman TJ, Day M, Perry N, Gally DL, Parkhill J, Jenkins C, Cain AK, Fraser CM, Collier RJ. 2018. Transposon Insertion Sequencing Elucidates Novel Gene Involvement in Susceptibility and Resistance to Phages T4 and T7 in Escherichia coli O157. mBio 9:e00705-18.

49. Rubin EJ, Akerley BJ, Novik VN, Lampe DJ, Husson RN, Mekalanos JJ. 1999. In vivo transposition ofmariner-based elements in enteric bacteria and mycobacteria. Proceedings of the National Academy of Sciences 96:1645–1650.

50. Kortright KE, Chan BK, Turner PE. 2020. High-throughput discovery of phage receptors using transposon insertion sequencing of bacteria. Proceedings of the National Academy of Sciences 117:18670–18679.

51. Dunstan RA, Bamert RS, Belousoff MJ, Short FL, Barlow CK, Pickard DJ, Wilksch JJ, Schittenhelm RB, Strugnell RA, Dougan G. 2021. Mechanistic Insights into the Capsule- Targeting Depolymerase from a Klebsiella pneumoniae Bacteriophage. Microbiology spectrum 9:e01023–21.

52. Pickard D, Kingsley RA, Hale C, Turner K, Sivaraman K, Wetter M, Langridge G, Dougan G. 2013. A genomewide mutagenesis screen identifies multiple genes contributing to Vi capsular expression in Salmonella enterica serovar Typhi. J Bacteriol 195:1320–6.

53. Gordillo Altamirano FL, Barr JJ. 2021. Unlocking the next generation of phage therapy: the key is in the receptors. Curr Opin Biotechnol 68:115–123.

54. Cota I, Sánchez-Romero MA, Hernández SB, Pucciarelli MG, García-Del Portillo F, Casadesús J. 2015. Epigenetic Control of Salmonella enterica O-Antigen Chain Length: A Tradeoff between Virulence and Bacteriophage Resistance. PLoS genetics 11:e1005667–e1005667.

55. Santander J, Robeson J. 2007. Phage-resistance of Salmonella enterica serovar Enteritidis and pathogenesis in Caenorhabditis elegans is mediated by the lipopolysaccharide. Electronic Journal of Biotechnology 10:627–632.

56. Guo G, Weiss B. 1998. Endonuclease V (nfi) mutant of Escherichia coli K-12. J Bacteriol 180:46–51.

