## Supplementary Information for "Collateral sensitivity increases the efficacy of a rationally designed bacteriophage combination to control *Salmonella enterica*"

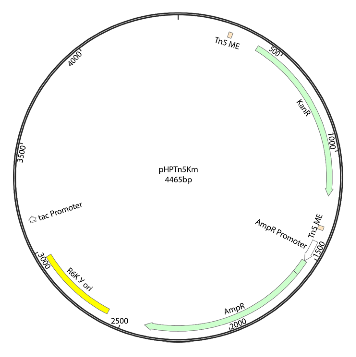

**Supplementary Figure 1. Genetic map of** plasmid pHPtn5Km. This plasmid was used to PCR amplify the Tn5::Km for preparation of transposomes used in the construction of a saturating transposon insertion mutant library.

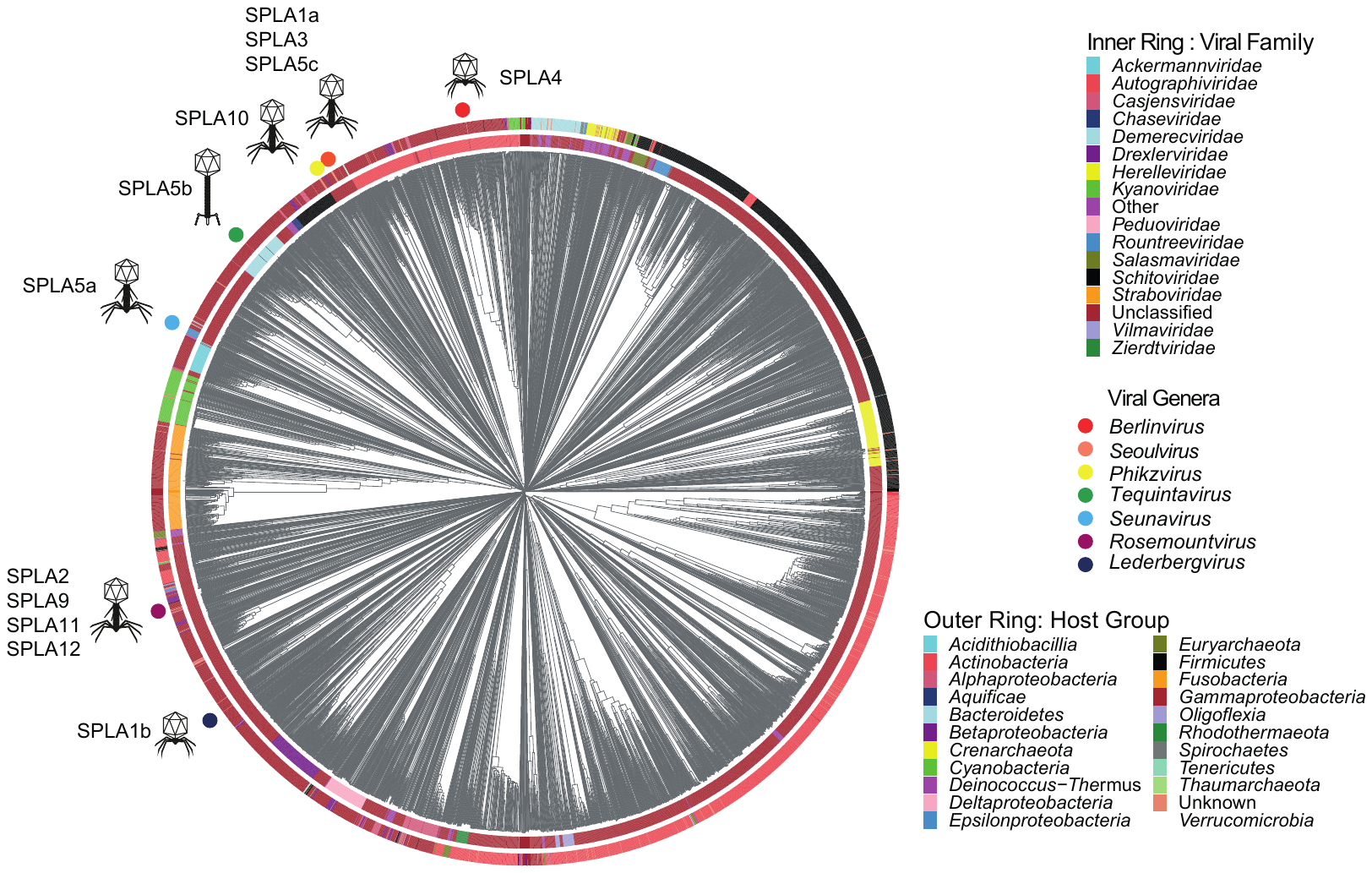

**Supplementary Figure 2. Relationship of SPLA phages in the context of viral proteomes from phages within the Viral-Host Database.** The dendogram and host group was generated using proteome data with VipTree and Viral Family was annotated using genomes from Inphared. SPLA Salmonella phage isolates coloured with circles for viral genera *Berlinvirus* (red), *Seoulvirus* (orange), *Phikzvirus* (yellow), Tequintavirus (green), *Seunavirus* (Lightblue), *Rosemountvirus* (Purple) and *Lederbergvirus* (darkblue). Icons indicate predicted phage morphology.

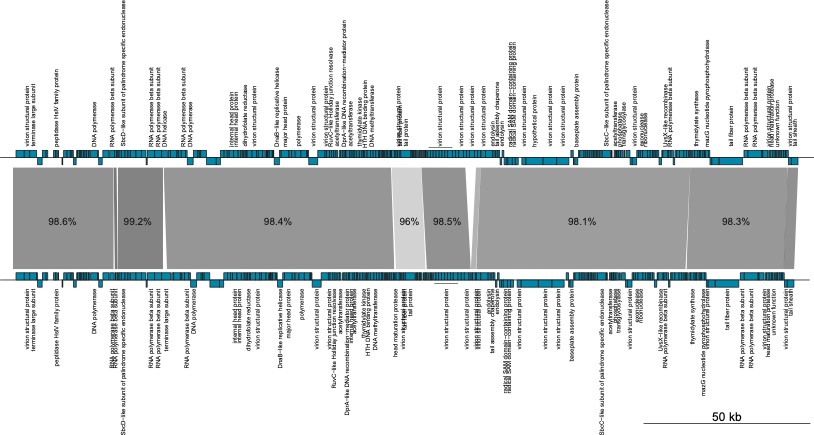

**Supplementary figure 3.** Whole genome alignment of *Seoulviruses* SPLA1a and SPLA5c with nucleotide sequence identity and predicted function annotated using PHROGS software. The intensity of the shading indicates the nucleotide sequence identity.

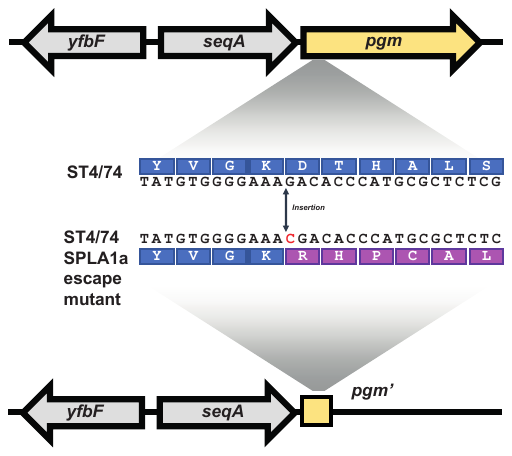

**Supplementary figure 4.** Nucleotide sequence alignment of *pgm* region of ST4/74 WT and ST4/74 SPL1a escape mutant (ST4/74*)

**Supplementary Tables available for download**

**Supplementary Table 1.** Bacterial strain collection used in the study for the isolation of bacteriophages and the investigation of bacteriophage host range.

**Supplementary Table 2.** List of Primer sequences used in the study.

**Supplementary Table 3.** List of log fold changes and statistical significance of transposon insertions within genes of Salmonella Typhimurium strain ST4/74 following exposure to six SPLA bacteriophages (SPLA1a, SPLA1b, SPLA2, SPLA5b, SPLA5c and SPLA11).
